# Viral isolation reveals novel and diverse phages infecting natural stream biofilms

**DOI:** 10.64898/2026.03.26.713887

**Authors:** Wai Hoe Chin, Martin Boutroux, Akira Harding, Davide Demurtas, Florian Baier, Hannes Peter

## Abstract

Bacteriophages of environmental bacteria remain underrepresented, lending paucity to phage-biofilm research beyond clinical and model species domains. Here, we present the **A**lpine **L**otic **P**hage (ALP) collection, curated through an isolation campaign from biofilm-forming bacteria of alpine streams. We obtained 57 phage isolates, which were dereplicated to 28 unique genomes following sequencing. The collection consists of tailed phages infecting 14 bacterial host species with genomes spanning 37 to 363 kb while exhibiting diverse plaque morphologies, depolymerase activity, and distinct impacts on host biofilm architecture. Comparative analyses against public viral genomes and a curated planetary-scale contig database revealed limited sequence similarity, underscoring the novelty of ALP phages. Functional annotation resolved 9 – 54% of predicted genes which encoded viral structural components, nucleotide metabolism functions, anti-defence mechanisms, and auxiliary genes that facilitate viral infection and replication. Together, the ALP collection represents a foundational resource for investigating phage evolution and ecology in natural bacterial communities.

## Introduction

Biofilms are ubiquitous microbial assemblages^1^, consisting of complex, spatially structured microbial communities that lend emergent properties such as resource retention and resilience against environmental stress^2^. These sessile communities have been under a negative spotlight in medical and industry settings but they play important roles in maintaining fundamental functions across a wide range of ecosystems. For example, benthic biofilms in streams and rivers orchestrate carbon and nutrient cycling at the base of lotic food webs^3,4^ while often acting as sentinels for monitoring climate-induced changes to the ecosystem^5,6^.

As the dominant microbial lifestyle for millienia^7^, biofilms also possess a long eco-evolutionary history with bacteriophages (phages for short), which are viruses that infect bacteria. Virulent phages adopt the lytic cycle exclusively where infected cells are lysed to release assembled virions, while temperate phages alternate between active lytic replication and dormant lysogeny as the phage genome integrates with the bacterial chromosome^8^. Through these lifecycles, phages regulate bacterial abundance and foster microbial diversity^9^, co-evolution, and horizontal gene transfer^10,11^. In doing so, phages are also the main source of microbial mortality in natural systems, affecting broader biogeochemical processes by liberating organic matter via cell lysis^12,13^ while also shifting metabolic demands of infected populations by reprogramming their hosts^14,15^. Phage predation is therefore expected to shape the ecology and evolution of biofilm communities^15^. However, the complex interactions between phages and biofilm-dwelling bacteria in nature remain poorly understood.

Major strides in unravelling phage-biofilm interactions were largely made within the motives of phage therapy research^16^. Key insights were derived from single-species biofilms of model and clinically-relevant bacterial species such as *Escherichia coli*, *Staphylococcus epidermidis*, and *Vibrio cholerae.* These studies have uncovered the protective effects of the biofilm matrix against phages^17^, the impact of cell packing density on viral spread across the biofilm^18^, and communal signalling amongst biofilm-dwellers under phage invasion^19,20^. Few have extended these insights to mixed-species biofilm^21,22^, yet these systems remain confined to model bacteria, thereby representing a narrow subset of phage-biofilm interactions. In contrast, field-based metagenomics of native biofilms have revealed robust coupling and host-specificity between phages and bacteria that extend to global scales^23,24^, often reporting microbial taxa that are underrepresented in contemporary biofilm research. However, a mechanistic basis underlying these field observations often remain unresolved due to a paucity of matched, culturable phage-bacteria pairs necessary for empirical investigation and validation. For instance, leveraging synthetic assemblages to mimic environmental biofilms and investigate viral-driven mechanisms in spatially structured heterogeneous communities. Many extensive and well-coordinated initiatives have been undertaken to develop and maintain phage repositories globally^25^ such as PhagesDB^26^, BASEL^27^, and CPL^28^. However, these collections remain largely focused on model and clinically relevant bacterial species with environmental strains remain comparatively underrepresented.

To advance the field beyond these confines, we present the **A**lpine **L**otic **P**hage (ALP) collection: a resource comprising of 28 unique phage isolates infecting 14 environmental biofilm-forming bacterial species, spanning three taxonomic classes (α- and γ-proteobacteria, and Flavobacteriia). By integrating morphological, phenotypic, and genomic characterization, the ALP collection represents a highly novel and diverse culturable fraction of viruses previously hidden within the viral “dark matter” of natural stream biofilms. We envisage this resource to provide a foundation for empirical and computational studies aimed at understanding phage–biofilm dynamics, adaptation, and ecosystem function within environmentally relevant contexts. This collection also offers a valuable reference for advancing phage gene annotation and exploring the biotechnological potential of alpine phages.

## Results

### Phage isolation workflow and overall composition of the ALP collection

To facilitate the isolation of phages infecting biofilm-forming bacteria in nature, we selected a panel of 37 bacterial isolates from a recently established in-house collection derived from stream biofilms. This panel spans 24 genera across eight taxonomic classes, representing bacterial taxa commonly reported in freshwater biofilm communities (Table S1). Notably, genera such as *Rhodoferax*, *Flavobacterium*, *Massilia*, and *Sphingomonas* are well-documented inhabitants of streambed biofilms, with *Rhodoferax* and *Flavobacterium* reported as abundant and widespread across glacier-fed streams globally^29,30^. These taxa contribute to key ecosystem processes, including carbon and nutrient cycling^24^, and are increasingly susceptible to climate-driven environmental change^6,31^.

All bacterial isolates (and their corresponding phages) grew on standard R2A media and encompassed a spectrum of phenotypes relevant to biofilms including a range of growth rates, surface motility, aggregative behaviour (i.e. floc-forming), and pigment production. Stream water was collected from the confluence of a groundwater-fed stream (La Vièze) and a glacier-fed tributary (La Saufla) (Fig.S1), which was subsequently concentrated using tangential flow filtration, followed by 0.45 μm dead-end filtration to remove most prokaryotic and eukaryotic cells while retaining the concentrated viral fraction (Fig. 1A). Phages were isolated using soft-agar overlays of the concentrated water samples where observed plaques were picked, double plaque-purified, and amplified for downstream characterisation by whole-genome sequencing and transmission electron microscopy (TEM) (Fig. 1A).

**Fig. 1.**
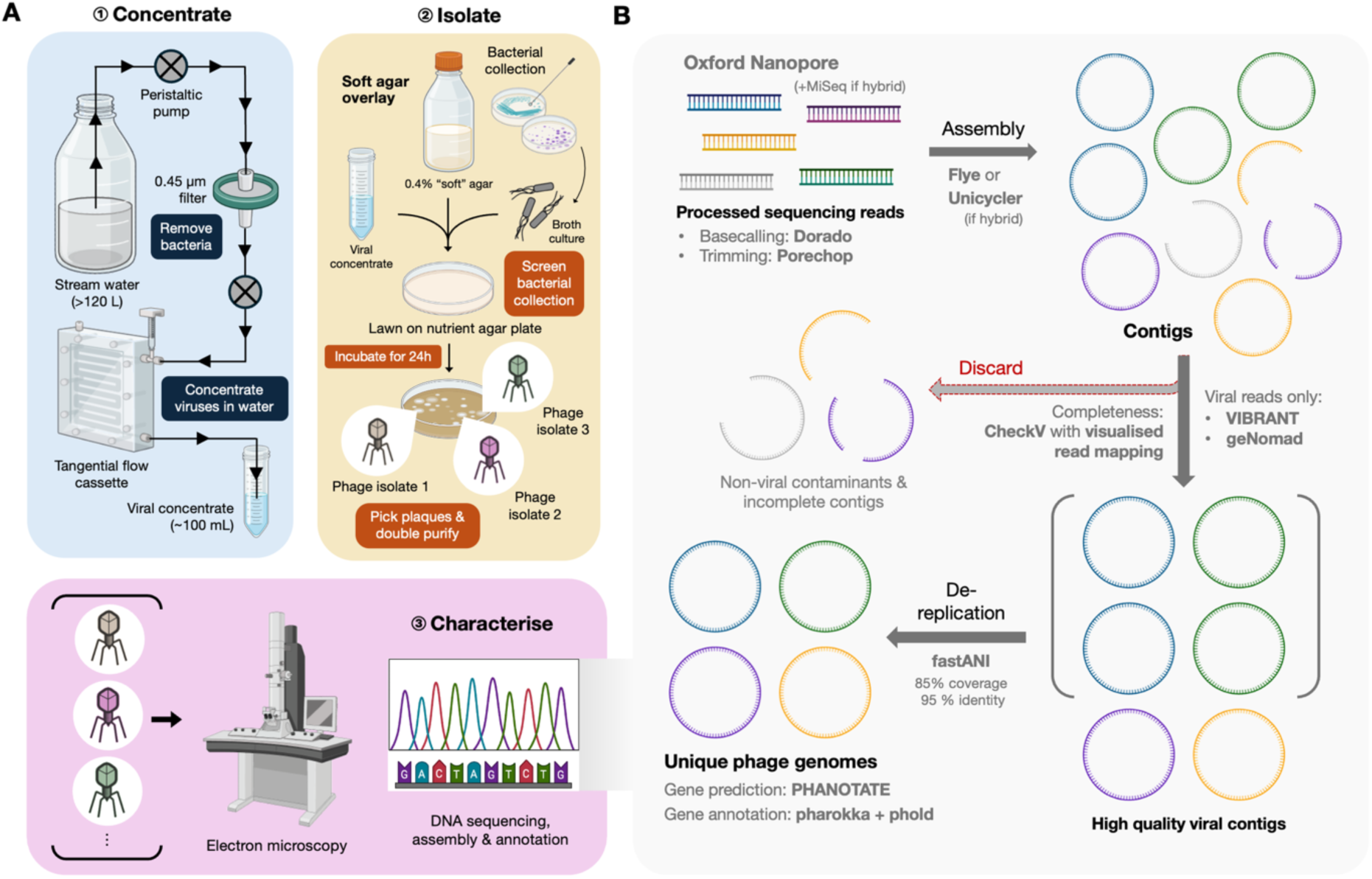
Phage isolation workflow and bioinformatic assembly pipeline of viral genomes. A) ∼120 L of stream water was collected from an alpine stream and was filtered to remove prokaryotes with a 0.45 µm filter. The viral fraction retained was concentrated by ∼100-fold via tangential flow. The concentrated stream water was screened for phages using soft-agar overlay mixed with bacterial host broth culture. Visible plaques were picked, double-purified, and amplified in host broth culture to yield pure high-titre phage lysates. These lysates were then leveraged to characterise phages via transmission electron microscopy (TEM) and whole-genome sequencing. B) Summary of DNA sequencing and bioinformatic pipeline to yield unique, high-quality, and complete phage genomes which constitutes the ALP collection. Hybrid assemblies with long-read Oxford Nanopore and short-read MiSeq sequencing platforms were adopted for selected isolates with marginal DNA quantity due to inherent challenges with DNA extraction. Tools adopted in each step of the pipeline are indicated by bolded grey text, where tool versions are referenced in online methods.

In total, 57 phage isolates were recovered across 14 bacterial host genera (Fig. 2A). *Rahnella inusitata* yielded the highest number of isolates (n = 22), followed by *Pseudomonas cyclaminis* (n = 7) and *Pseudomonas haemolytica* (n = 6). All isolates were culturable except a single *Sphingomonas* phage, which could not be amplified sufficiently for DNA extraction and transmission electron microscopy (TEM). High-quality genomic DNA was obtained from 43 of the 56 culturable isolates, whereas the remaining isolates - particularly those infecting *P. cyclaminis* - were recalcitrant to DNA extraction. Dereplication based on genome identity resulted in a final collection of 28 unique phages (Fig. 2A; Table S2). All 28 assembled genomes were independently classified as viral by both VIBRANT and geNomad. CheckV assessment indicated that 26 genomes were high quality and 2 (Rahnella phages B311P5 and B311P9) were medium quality (Table S3). All genomes were assigned by geNomad to the class *Caudoviricetes*, with eight further resolved at the family level: six *Autographiviridae*, one *Casjensviridae*, and one *Schitoviridae* (Table S3). Across host genera, between 1 and 9 unique phages were recovered per bacterial genus. Notably, the 22 initial Rahnella-infecting isolates dereplicated into nine distinct phages, indicating frequent re-isolation of closely related viruses targeting this host (Fig. 2A).

**Fig. 2.**
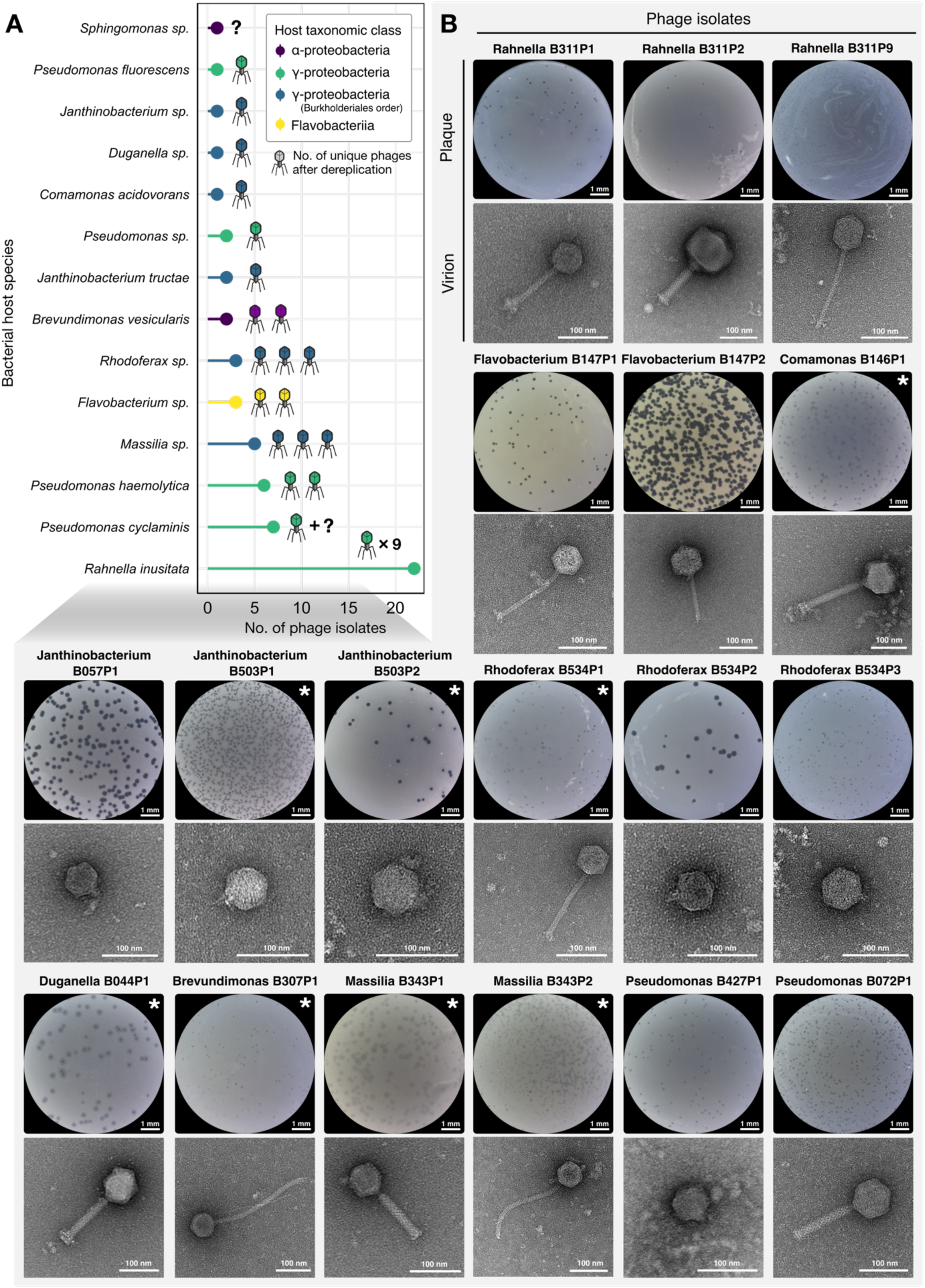
Plaque and virus morphologies of alpine stream phages. A) A total of 28 unique phage isolates were derived from 14 alpine bacterial species in our viral screening effort. Coloured legend represents the bacterial host taxonomic class that were targeted by phages while phage icons represent the number of genomes after dereplication (e.g. 2 unique phages were isolated for *Flavobacterium* sp.) that were isolated per bacterial species. Question mark designates the unknown number of unique phages as dereplication was not possible without successful phage genome extraction (except *Sphingomonas* sp. phage which only had one isolate but DNA extraction was unsuccessful). B) Phage plaques were assayed using 0.4% soft agar overlay with brightfield images demonstrating a variety of plaque sizes and morphologies. Haloed plaques demonstrating potential depolymerase activity were marked with asterisks. Transmission electron microscopy images of 18 phage isolates revealed a collection of tailed phages consisting of myoviruses, podoviruses, and siphoviruses. Only the *Sphingomonas* phage isolate was neither successfully sequenced nor imaged due to poor culturability *in vitro*.

### Alpine stream phages exhibit diverse viral and plaque morphologies

Unique viral isolates were imaged with TEM to provide a first glance at the morphotypes pertaining to phages infecting stream biofilms. Of the 28 phage isolates, we were able to acquire electron micrographs for 18 (Table S2). TEM images revealed a collection of tailed viruses comprising 7 siphoviruses, 6 podoviruses, and 5 myoviruses, highlighting the heterogeneous viral population infecting sessile bacterial communities in streams (Fig. 2B). Two myoviruses, Rahnella phage B311P2 and Comamonas phage B146P1, exhibited large virion dimensions with capsid diameters exceeding 100 nm, consistent with structural dimensions expected of jumbophages^32^.

In addition to virion structure, plaque morphologies were assessed for the 18 imaged isolates to capture basic phage phenotypes under semi-solid growth conditions. Here, plaque sizes varied markedly across the collection, ranging from small punctate plaques to large clearings up to 280 μm in diameter (Fig. 2B). Plaques exceeding 200 µm in diameter were predominantly produced by podoviruses, in line with their smaller virion dimensions which enhances diffusivity in structured matrices relative to larger myo- and siphoviruses. Recent biophysical insights further suggest that larger plaque sizes may also be associated with shorter viral latent periods which lends higher phage virulence^33^. Plaques also differed in qualitative features, including central turbidity and translucent halos extending beyond the plaque margin. Most isolates produced clear plaques reflecting their effective host-killing capacity. However, Massilia phage B343P1 and Pseudomonas phage B427P1 formed turbid plaques, indicative of reduced lytic capacity relative to other ALP isolates.

In addition, 8 of the 18 imaged phages produced translucent halos surrounding the plaque border, with radii ranging from 30 to 250 µm (Fig. 2B). These halos are consistent with depolymerase activity, whereby phage-encoded enzymes degrade bacterial surface polysaccharides including biofilm exopolysaccharides to facilitate access to host receptors^34^. Intriguingly, Janthinobacterium phages B503P1 and B503P2 were dereplicated as near-identical genomes yet, the isolates exhibited distinct plaque morphologies (Fig. 2B). Isolate B503P2 produced larger plaques with more extensive depolymerase halos compared to B503P1, indicating phenotypic divergence despite 99.9% genome identity. Closer inspection revealed a single nucleotide substitution in the tail fibre gene, resulting in a serine-to-asparagine substitution at position 590 (S590N; Fig.S2). Although the structural consequences of this substitution is not known, it occurs adjacent to a region partially homologous to a pectate lyase domain, which was associated with depolymerase activity in *Acinetobacter* phages^35^. Collectively, these observations reveal the substantial morphological and phenotypic diversity of lotic phages where the latter underscores the various viral infection traits leveraged against structured sessile microbial communities.

### The ALP collection represents an emerging repository of novel viruses infecting stream biofilm-forming bacteria

Given the understudied nature of viruses associated with natural stream biofilms, we assessed the genomic novelty represented by the ALP collection. Briefly, phage genomes were BLAST-queried against the NCBI nucleotide database where we retained the top five hits based on sequence coverage and nucleotide identity. The majority of isolates exhibited limited similarity to previously described viruses, with <20% genome coverage and a mean nucleotide identity of ∼80% (Table S4). Only 12 of the 28 phages showed >20% coverage against publicly available viral genomes, indicating that most ALP phages lack close relatives in existing reference databases and public collections (Table 1). Among these, Pseudomonas phage B508P1 demonstrated the strongest match (94% genome coverage and nucleotide identity) with a cold-active podovirus VSW-3 infecting *Pseudomonas fluorescens* in wetlands^36^. This is followed by Rahnella phage B311P4 with 96% coverage and 87% identity against phage KLB24 infecting plant-associated *Klebsiella* sp. which was isolated from farmland rain puddles^37^. Further adding to the relatedness of ALP isolates with other aquatic phages, Pseudomonas phage B072P2 displayed 81–86% genome coverage and ∼85% nucleotide identity to podoviruses also infecting *P. fluorescens*, notably the psychrophilic freshwater phage phiGM22-3^38^. The remaining phages infecting *Rahnella*, *Massilia, Brevundimonas*, and *Rhodoferax* exhibited varying similarity (23–50% coverage, >74% identity) to known phages, except Rahnella phage B311P3 which was partially found amongst *Serratia* bacterial genomes, implying a possible distant prophage relative within this genus. We also note that Rahnella jumbophage B311P2 appears distantly related to two other jumbophages Cronobacter phage vB_CsaM_GAP32 and Escherichia phage PBECO 4, suggesting that these large viruses could share core genomic segments.

**Table 1:** List of top two BLAST with >20% query coverage of ALP phages against NCBI database.

| Phage (φ) isolate | Closest hits | Query coverage | % identity | Accession # |
| --- | --- | --- | --- | --- |
| Brevundimonas φ B307P1 | *Caudoviricetes* sp. (MAG)<sup>a</sup> | 38% | 80.94% | OR222513.1 |
|  | *Caulobacter* phage Seuss | 14% | 77.03% | NC_047757.1 |
| Brevundimonas φ B307P2 | *Caudoviricetes* sp. (MAG)<sup>a</sup> | 36% | 81.23% | OR222513.1 |
|  | *Caulobacter* phage Seuss | 14% | 77.02% | NC_047757.1 |
| Massilia φ B343P2 | *Caudoviricetes* sp. (partial MAG)<sup>a</sup> | 68% | 85.50% | BK020520.1 |
|  | *Caudoviricetes* sp. (MAG)<sup>a</sup> | 1% | 75.35% | OR222459.1 |
| Pseudomonas φ B072P2 | *Pseudomonas* phage phiGM22-3 | 86% | 84.73% | MW627366.1 |
|  | *Pseudomonas* phage phi2 | 81% | 85.57% | NC_013638.1 |
| Pseudomonas φ B508P1 | *Pseudomonas* phage VSW-3 | 94% | 94.39% | NC_041885.1 |
|  | *Providencia* phage PSTNGR1 | 2% | 74.38% | MW145136.1 |
| Pseudomonas φ B529P1 | *Pseudomonas* phage phiPsa300 | 50% | 82.01% | NC_073687.1 |
|  | Pseudomonas phage phiPsa347 | 49% | 81.99% | NC_073685.1 |
| Rahnella φ B311P2 | Cronobacter phage vB_CsaM_GAP32 | 33% | 72.00% | JN882285 |
|  | Escherichia phage PBECO 4 | 15% | 68.00% | KC295538 |
| Rahnella φ B311P3 | *Serratia fonticola*<sup>b</sup> | 23% | 79.02% | CP013913.1 |
|  | *Serratia* sp. JSRIV004<sup>b</sup> | 22% | 79.07% | CP074140.1 |
| Rahnella φ B311P4 | Bacteriophage sp. (partial MAG)<sup>a</sup> | 96% | 87.22% | OP072654.1 |
|  | Klebsiella phage vB_KM5a1-KLB24 | 96% | 87.13% | PP554338.1 |
| Rahnella φ B311P5 | Enterobacter phage EC151 | 27% | 74.50% | MW464860.1 |
|  | Klebsiella phage vB_Ko_K4PH164 | 8% | 73.92% | OY979482.1 |
| Rahnella φ B311P7 | Enterobacter phage EC151 | 36% | 76.76% | MW464860.1 |
|  | Klebsiella phage vB_Ko_K4PH164 | 22% | 75.07% | OY979482.1 |
| Rhodoferax φ B534P2 | Variovorax phage Gard | 38% | 80.25% | PV920657.1 |
|  | Variovorax phage VAC_51 | 32% | 79.67% | OX359471.1 |

<sup>a</sup> MAG denotes metagenome-assembled genomes that are either complete or partial.
<sup>b</sup> Closest hits with bacterial origins i.e. non-viral hits.

To further contextualise novelty at a global scale, the ALP genomes were queried against LOGAN, which is a planetary-scale contig database curated from public sequence read archives^39^. Using a permissive k-mer coverage threshold of 0.25, 16 out of 28 ALP phages returned positive matches (Fig. 3A). These matches were geographically widespread, predominantly originating from the European continent but also spanning Asia and North America (Fig. 3A, Table S5). The matches were also associated with diverse environmental sources ranging from freshwater ecosystems including rivers, glaciers, groundwater and alpine streams to other environments such as soil, plants, and terrestrial animals, followed by anthropogenic food and wastewater sources (Figs.3A and B). Overall, 7 phage isolates matched with contigs from a single source while the remaining 9 isolates were associated with contigs from multiple environments of up to 6 sources (Fig. 3B). Despite this broad distribution, individual matches generally exhibited low similarity, with a median k-mer coverage score of 0.36 (range: 0.25–0.85) against LOGAN contigs (Fig. 3A). Collectively, the sparse representation of phages in both reference and global metagenomic databases underscores the substantial novelty behind viruses associated with stream biofilms.

**Fig. 3.**
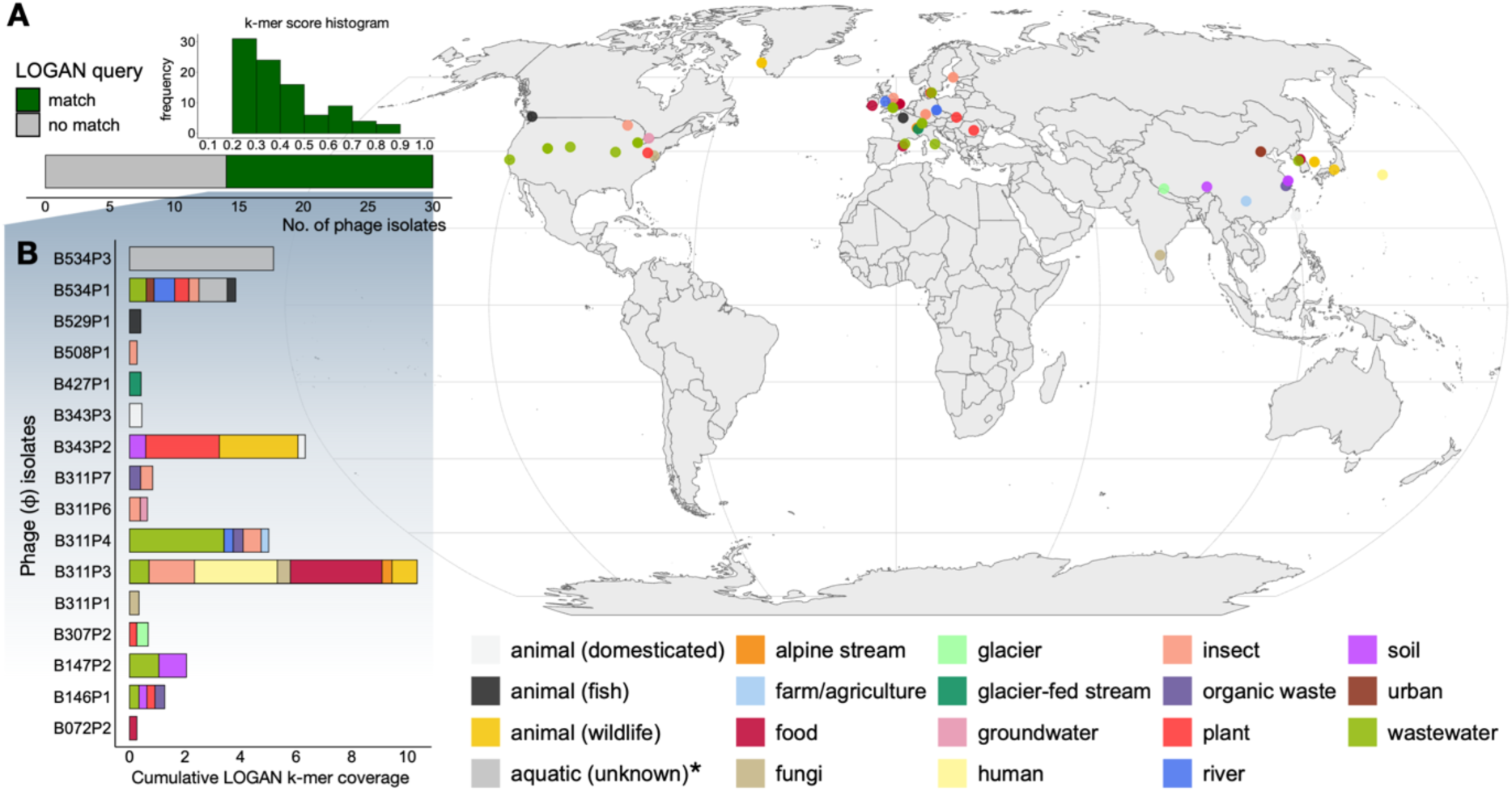
Matches of ALP genome sequences against planetary-wide contig database. A) Total number of phage isolates with matches to contigs within the LOGAN database and a histogram depicting the spread of k-mer coverage scores of all positive hits to the database. The map represents the geographical distribution of these hits while the colours represent the source of the contigs identified within the database. Asterisk denote matches to “aquatic (unknown)” sources do not possess any geographical metadata and are thus, not located on the map. B) Cumulative k-mer coverage of sequence matches between a 1kb sequence of the ALP isolates to the LOGAN contigs, where colours also represent the contig source stipulated within the database.

### Auxiliary coding sequences in phages facilitate host takeover for infection and environmental resilience

Across the 28 phage isolates, genome sizes spanned over an order of magnitude from 37 to 363 kb (Fig. 4A). Two isolates, Comamonas phage B146P1 (206 kb) and Rahnella phage B311P2 (363 kb), exceeded 200 kb and were hence, classified as jumbophages consistent with their large virion dimensions (Fig. 2B). Differences in phage genome size were matched by expansions in capsid size, with phages below 83 kb exhibiting relatively similar capsid diameters (69 nm on average). Meanwhile, capsids enlarged progressively as genome size exceed 111 kb, reaching 133 nm in the Rahnella jumbophage B311P2 with a 363 kb genome. Allometric analysis also revealed proportional scaling between genome size and estimated capsid volume (allometric coefficient α = 0.91), indicating that DNA packing density was relatively conserved across the ALP collection following structural principles of icosahedral dsDNA viral capsids^40^ (Fig. 4B, Table S6). Furthermore, coverage patterns from read mapping identified DNA packaging mechanisms^41^ for 14 phage isolates: 1 utilized a 3’ *cos* terminus, 4 harboured *pac* sites, and 9 employed direct terminal repeats (DTR) (Table S7), where the Rahnella jumbophage B311P2 possessed a remarkably long DTR of 19,824 bp. Together, these features illustrate the genomic architectural diversity, packaging constraints, and mechanisms among isolates of the ALP collection.

**Fig. 4.**
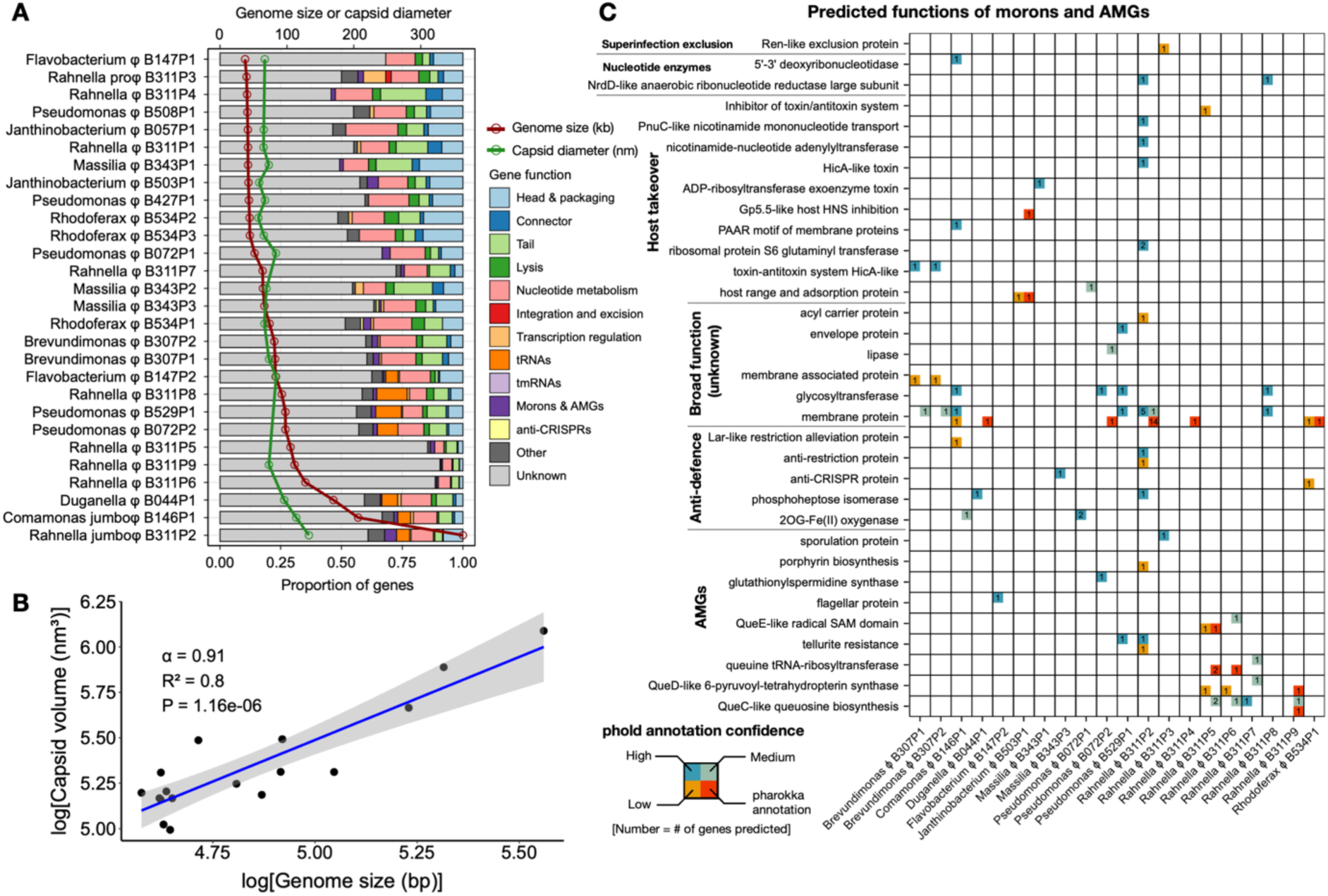
Genomic features of ALP isolates and profile of auxiliary coding sequences. A) Genome analysis of sequenced phage isolates where the red and green line plots correspond to genome size (kb) and capsid diameter (nm) respectively, which are plotted on the top x-axis, while the coloured bar chart indicates the proportion of genes predicted, corresponding to the bottom x-axis. B) Log-transformed plot of capsid volume (approximated as spherical volume: [ν ÷ 6] × *D*^3^, where *D* is the capsid diameter) against genome length. The regression line is shown in blue with the accompanying allometric exponent (a i.e. gradient of the log-transformed regression), R-squared value, and p-value. C) Frequency table of morons and AMGs predicted in 20 phage isolates using nucleotide-based (pharokka) and protein structure-based (phold) algorithms. Gene functions are broadly categorised into 6 groups. Each cell within the table is separated to four quadrants which are colour-coded based on phold annotation confidence i.e. (blue = high, green = medium, and low = orange), while the bottom-right red quadrant indicates genes predicted by the nucleotide-based algorithm, pharokka. The number within each quadrant shows the frequency of gene functions predicted in accordance with the type of algorithm employed and confidence level for phold annotations.

Genome annotation resolved putative functions for 9 – 54% of predicted coding sequences (CDS), with gene functions predominantly associated with virion structure, nucleotide metabolism, and host lysis (Fig. 4A, Table S6). Only one isolate – Rahnella phage B311P3 – was identified as a temperate phage, encoding both integrase and excisionase while also possessing genes consistent with lysogenic capacity such as CII- and CIII-like transcriptional regulators and Ren-like superinfection exclusion factor. In addition, Rahnella phage B311P3 was also detected as an integrated prophage within the *Rahnella* host genome. Meanwhile, larger phages generally encoded substantial tRNAs relative to the entire collection, comprising 5 – 11% of total CDS (Fig. 4A). This pattern aligns with previous observations that phages with larger genomes leverage self-proprietary tRNAs to sustain viral replication as host translational capacity becomes limiting during late-stage infection^42^.

Beyond core functions, morons and auxiliary metabolic genes (AMGs) were predicted in 20 of 28 phage genomes, ranging from 1 – 8 genes per genome except for the Rahnella jumbophage, which encoded 33 auxiliary genes (Figs.4A and C, Table S8). The majority of these genes however, corresponded to membrane proteins and biocatalytic enzymes of broad unclassified functions. Among fully annotated genes, functions associated with host takeover and anti-defence were prevalent, such as the host mRNA-modulating HicA-like toxin-antitoxin system^43^, NAD synthesis and scavenging factors, followed by anti-restriction and anti-CRISPR systems. Several phages also encoded enzymes implicated in phage DNA modification, such as 2-oxoglutarate/Fe(II)-dependent oxygenases^44^ and phosphoheptose isomerases^45^, consistent with strategies to evade intracellular host restriction and CRISPR-Cas immunity (Fig. 4C).

In contrast, relatively few AMGs were predicted to directly enhance host environmental resilience. These included tellurite resistance genes encoded by the Rahnella jumbophage and Pseudomonas phage B529P1, as well as genes involved in antioxidative protection and cellular energetics in other isolates (e.g. gluthathionylspermidine synthase and porphyrin biosynthesis; Fig. 4C). These AMGs are consistent with promoting survival and maintaining cellular metabolism in biofilm-associated microenvironments characterised by accumulation of trace metal ions^46,47^, steep redox gradients^48^, and chronic oxidative stress^49^. Notably, 4 Rahnella phage isolates encoded enzymes within queuosine biosynthesis pathway alongside a queuine tRNA-ribosyltransferase, suggesting roles in phage DNA modification for host immune evasion^50^ and maintaining translational efficiency to sustain bacterial cells in biofilms^51^. Despite expectations that temperate phages preferentially retain host-beneficial genes^52^, only a single AMG was detected in the temperate Rahnella phage B311P3, encoding a sporulation factor of unclear relevance to the non-sporulating *Rahnella* genus. Collectively, these findings reveal that stream-biofilm phages encode diverse auxiliary gene functions primarily associated with host takeover and antiviral defence evasion, with a smaller subset linked to host cellular and biofilm stress tolerance. The large unannotated portion of ALP genomes also represents a resource for exploring additional phage-encoded adaptations or even, gene products of potential biotechnological relevance.

### Phage infections in microfluidics reveal host-specific architectural responses of environmental isolate biofilms

Microfluidic platforms have emerged as powerful systems to resolve phage-biofilm interactions with high spatial and temporal precision, lending crucial insights into mechanistic responses of biofilms under phage infection^17,20,21,53^ and viral propagation strategies in structured communities^18,54^. Yet, their application to environmentally derived phage-host systems remains limited. To demonstrate the utility of the ALP collection, we performed time-course infections of environmental isolate biofilms in microfluidic devices. We selected three representative bacterial isolates exhibiting high, moderate, and low biofilm-forming capacities (*Massilia* sp., *R. inusitata*, and *P. fluorescens*, respectively), based on crystal violet staining of static overnight biofilms (Fig. 5A). These isolates were seeded into microfluidic channels and allowed to establish for 24 hours under continuous flow prior to infection with their corresponding phages: Massilia phage B343P1, Rahnella jumbophage B311P2, and Pseudomonas phage B508P1 (Fig. 5B).

**Fig. 5.**
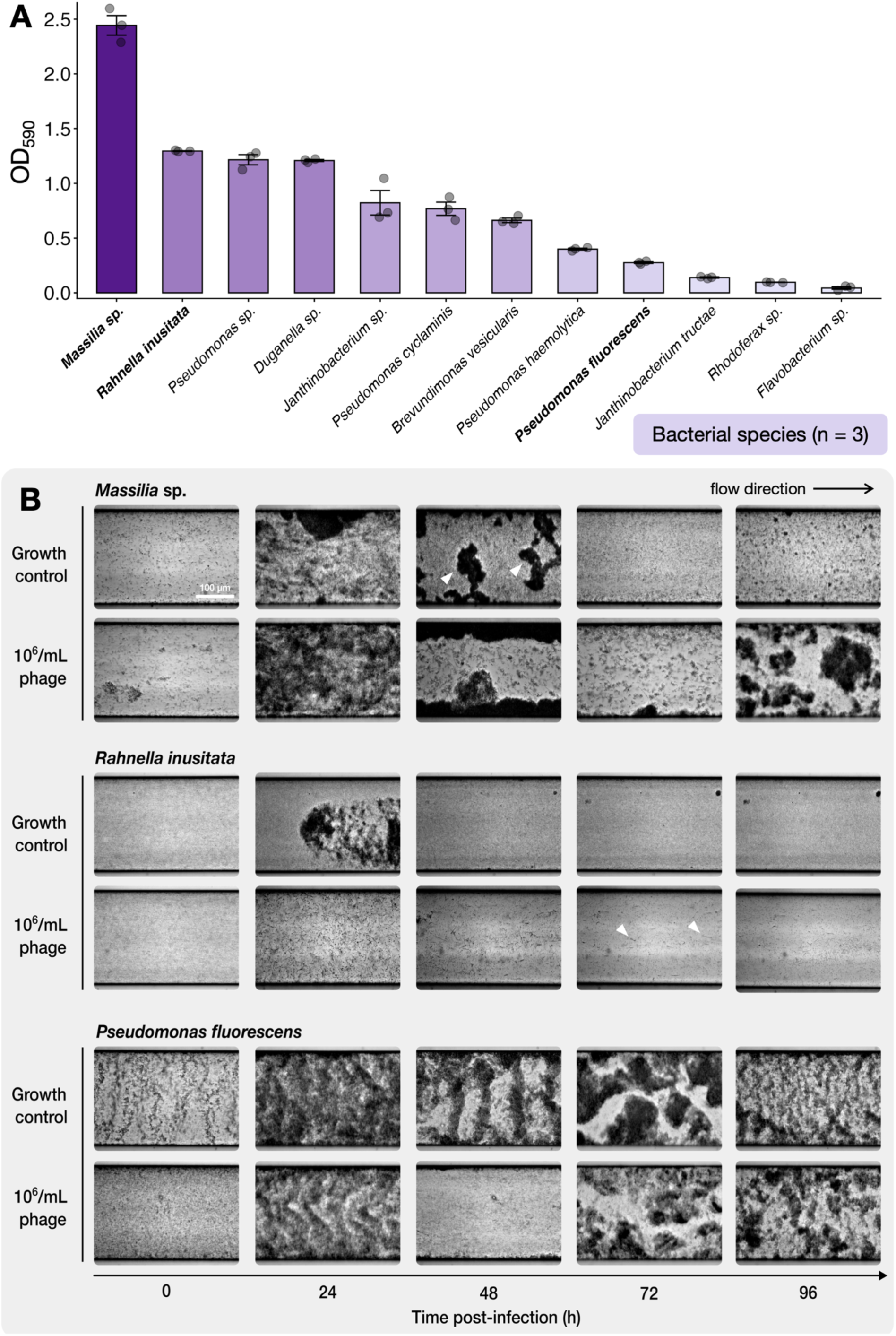
Biofilm forming capacity of environmental isolates and time-course phage infection in microfluidics. A) Crystal violet absorbance A_590_ values for static biofilms grown in 96-well plates for 24 hours with three replicates (n = 3) per bacterial species. Species highlighted in bold were adopted for microfluidic experiments as representative for their varying capacities to form biofilm (i.e. high, moderate, and low). Error bars represent the SEM of triplicate A_590_ values. B) Brightfield images of 24 h-old biofilms established in microfluidic devices which were subsequently infected with 10^6^ phages/mL under continuous flow at 0.1 µL/min and monitored up to 96 hours post-infection. *Massilia* sp. biofilms were infected with phage B343P1, *R. inusitata* with jumbophage B311P2. and *P. fluorescens* with phage B508P1. Arrowheads in *Massilia* sp. biofilm highlights examples of microcolonies observed over the experimental duration while arrowheads in *R. inusitata* community indicate cell filamentation under sustained jumbophage pressure.

Over the course of 96 hours following infection, biofilms were monitored using brightfield microscopy. These environmental isolates are non-fluorescent and display heterogeneous growth patterns under fluid flow (Fig. 5B), highlighting the exploratory potential of extending microfluidic phage-biofilm assays beyond fluorescently tagged model systems. Here, time-course imaging revealed distinct architectural characteristics across the three biofilm species. *Massilia* sp., a floc-forming isolate, developed heterogeneous biofilms with microcolonies reaching up to ∼200 µm wide. Meanwhile, environmental *P. fluorescens*, despite limited biofilm formation in static assays, formed dense communities with patterned fronts that aligned with the direction of flow. In contrast, *R. inusitata* formed comparatively homogeneous and low-density lawn-like biofilms spanning the channel surface (Fig. 5B). Notably, *Massilia* sp. and *P. fluorescens* biofilms exhibited quasi-cyclical development even in the absence of phage, characterized by initial disruption of heterogeneous biofilms under flow between 48 – 72 hours followed by structured re-growth.

Phage predation reduced biofilm surface area across all systems by 48 h post-infection and altered the structure of *Massilia* sp. and *P. fluorescens* communities. Despite continuous viral pressure, *Massilia* sp. and *P. fluorescens* biofilms recovered but adopted architectures distinct from both their early developmental states and uninfected controls. *Massilia* sp. biofilms formed sparser microcolonies, whereas the structured fronts of *P. fluorescens* appeared distorted despite unchanged flow conditions (Fig. 5B, 96 hpi). These changes imply potential trade-offs between biofilm formation and phage resistance in these two isolates. In contrast, *R. inusitata* biofilms failed to recover under sustained jumbophage pressure, and surviving cells frequently exhibited filamentation similar to infections caused by *E. coli* jumbophage Sharanji^55^. Together, these findings demonstrate that phages can actively reshape biofilm architecture and induce host-specific responses that dictate biofilm recovery under sustained viral pressure. By extending microfluidic infection assays beyond model systems, these experiments illustrate the tractability of environmentally derived phage-host pairs within contemporary biofilm platforms and provides a framework for investigating virus-biofilm dynamics in structured natural communities.

## Discussion

Biofilms represent the dominant microbial lifestyle on Earth, forming highly structured and ubiquitous communities that underpin essential ecosystem processes^1^. As obligate bacterial parasites, phages have coexisted with biofilms over evolutionary timescales, and metagenomic surveys consistently reveal strong coupling between viral and bacterial community composition across diverse environments^8,23,24,56^. Despite the central role of phages in driving microbial diversity, carbon cycling, and horizontal gene transfer^57^, our understanding of phage-biofilm interactions remains disproportionately informed by model and applied phage-host systems. Consequently, the diversity, biology, and ecological roles of phages in natural multispecies biofilms remain poorly understood. The ALP collection addresses these gaps by providing a culturable, full-genome resource of novel, environmentally derived phages that extend our current understanding of viral diversity and traits beyond model and clinical biofilms.

Mountain stream ecosystems are structured and sustained by benthic biofilms that drive primary production, respiration, and nutrient cycling^3^. Through these processes, they support the functioning of river networks that supply freshwater to meet the demands of millions worldwide^58^. Moreover, stream biofilms are also sensitive indicators of environmental change and are increasingly affected by climate warming^6,29,31^. Therefore, there is growing urgency to understand how biofilms and their associated viral communities respond with accelerating climatic pressures. By targeting native stream bacteria from benthic biofilms, the ALP collection captures a fraction of viral diversity that is largely absent from existing phage and genome repositories. Although derived from alpine stream systems, its ecological relevance extends broadly as benthic biofilms are pervasive across aquatic ecosystems.

The limited representation of freshwater biofilm-associated phages in reference databases and sequence archives underscores the novelty observed across the ALP collection. Only 5 of 28 isolates exhibited >50% genome coverage with previously described freshwater phages (Table 1), while queries against the curated LOGAN database revealed sparse and low-coverage matches across geographically diverse environments (Fig. 3). The latter also indicates that potential relatives – or perhaps the ALP isolates themselves – are rarely recovered as near-complete genomes in metagenomic datasets, reflecting persistent technical challenges in viral metagenomic assemblies^59,60^. In contrast, DNA from purified isolates are more amenable to yield complete genomes thereby, providing greater confidence in identifying features such as terminal repeats that may inform phage DNA packaging strategies^41^. Moreover, the ALP isolates were sequenced and characterised with minimal laboratory passage, providing a complementary perspective of genomic content that reflect their recent life history in nature and minimising genetic drift caused by extended propagation under laboratory conditions^61^. In these contexts, the ALP collection not only expands taxonomic representation but also offers high-resolution genomes for facilitating bioinformatic analyses and ecological interpretation of biofilm-associated viruses in freshwaters.

Consistent with the global predominance of tailed dsDNA viruses^24,62^, the ALP collection comprises myo-, sipho-, and podovirus morphotypes supported by TEM imaging and genome annotation (Figs.2B and 4A). Capsid-genome scaling of the ALP isolates also followed allometric relationships consistent with icosahedral dsDNA phages, where scaling models reported proportional expansion in capsid volume with increasing viral genome sizes^40,63^ (Fig. 4B). This ensures that DNA packing density is maintained to preserve the internal capsid pressure required for genome ejection^40,63^, thereby reflecting the conserved infection mechanobiology amongst ALP isolates. Consequently, the ALP collection also expands the empirical basis of these capsid-genome scaling models^63^ by contributing physical measurements of virions derived from these novel isolates. Importantly, the conserved capsid-genome allometry also underscores the constraints of phages operating within environmental biofilms. For instance, larger capsids are expected to reduce phage diffusivity in biofilm matrices^64^, yet simultaneously accommodate larger genomes that may encode beneficial auxiliary genes, such as those enhancing host metabolism to support viral replication. While not directly tested here, this hints that biofilm-associated phages likely occupy a constrained multidimensional trait space, maintaining internal capsid pressure while balancing gene content, and diffusion through biofilms. Whether matrix-degrading depolymerases can partially alleviate diffusional constraints also remains unclear. This framework can be extended to stream environments where hydraulic forces (e.g. turbulent flow and shear) and oligotrophy may further shape viral structural stability and auxiliary gene content.

The compact and densely packed nature of phage genomes imposes strong constraints on gene content, which limits retention of genes beyond core replicatory functions^65^. Accordingly, morons and AMGs were sparse among ALP isolates and were largely associated with host takeover, metabolic reprogramming, and suppression of antiviral defences (Figs.4A and 4C). This pattern aligns with the predominance of virulent phages in the collection, prioritising genes that optimise the lytic cycle which typically occur at shorter timescales than lysogeny (Fig. 4C). In contrast, only a minor subset of AMGs was putatively linked to longer-term host environmental resilience, notably metalloid resistance and oxidative stress mitigation. These functions are congruent with overcoming stream biofilm-associated stressors, where EPS matrices are known to concentrate heavy metals and resident cells experiencing chronic oxidative stress^46,48^. Within the context of virulent phages, these AMGs may enhance phage production indirectly by temporarily sustaining host physiology under environmental stress rather than promoting long-term host persistence. Nonetheless, the true functional diversity of the ALP collection is likely underestimated given the overwhelming proportion of unknown gene functions (Fig. 4A), despite recent major strides with protein structure-based annotations in revealing the functions of >50% of genes within an average phage genome^66^.

Beyond genomic features, the presence of multiple infection phenotypes especially amongst phages infecting the same species (Fig. 2B; *Rhodoferax* and *Massilia* phages), coupled with host-specific structural responses to phage infection (Fig. 5B) suggests that viral invasion of biofilms is unlikely dictated by a dominant infection strategy. Instead, competing viral traits and unique host-specific adaptations in biofilms may stabilise microbial and viral diversity^56^ by promoting localised negative frequency-dependent “kill-the-winner” dynamics and limiting selective sweeps across the structured community. These effects are also likely amplified within natural multispecies biofilms as several taxa are competing within a spatially heterogenous niche, generating a mosaic of localised virus-microbe interactions where viral suppression is tempered and bacterial adaptation is dependent on the microenvironment^67^ and local phage pressure. In this context, environmental phages function not only as agents of mortality but as active architects of biofilm structure (Fig. 5B) and community composition, promoting turnover and preserving functional resilience of environmental biofilm by maintaining microbial community diversity.

In summary, the ALP collection captures substantial genotypic and phenotypic diversity among biofilm-associated phages in streams. This curated set of 28 unique phage isolates expands upon the viral “dark matter” that remains poorly represented in current culture collections and databases, particularly among underexplored natural environments where biofilms are pervasive. Crucially, the collection provides a foundation for necessary empirical investigations into virus-microbe interactions across scales. For example, integrating the ALP collection with synthetic communities^68,69^ and microfluidic platforms^18,21^ offers a tractable framework to finely dissect viral-driven mechanisms in heterogeneous biofilms. These experiments are also extendable to mesoscales such as flume models^70^ or interconnected fluidic systems of biofilm networks^71^ to explore and validate emergent ecological properties and evolutionary dynamics as phages propagate across broader biofilm landscapes. Beyond experimental applications, the viral genomes offer valuable references for advancing virome research and computational tools, with the prospect of uncovering environmental gene functions that may provide the biotechnological innovation required to navigate our rapidly changing ecosystems in a warming planet.

## Materials and methods

### Bacterial host and phage culture

All bacterial host isolates used in this study were cultivated at ambient room temperature (∼22°C) in 1× Reasoner’s 2a (R2A) medium (Neogen, USA) which was prepared following manufacturer’s protocol. Agar (Merck, Germany) was added to a final concentration of 0.4% or 1.5% for soft-agar overlay assay and standard agar media plates, respectively. Phages were propagated via broth or soft-agar medium depending on the viral isolate. Broth amplification was started with a 1:50 dilution of an overnight host culture in 10 mL of fresh media broth followed by spiking with 20 µL of phage lysate (minimum ∼10^6^ PFU/mL). The mixture is then incubated at room temperature with agitation overnight or until complete lysis i.e. broth clearance. The amplification reaction is centrifuged at 5000 ×g for 10 minutes to pellet bacterial debris and the viral supernatant is sterilised with 0.45 µm syringe filters (Sartorius, Germany). Amplification via soft-agar medium begins with mixing 500 µL of overnight host culture with 50 µL of serially-diluted phage lysate and 2 mL of 0.4% R2A agar. The 0.4% mixture is casted on 1.5% agar media to form a soft-agar overlay and the plate is incubated at room temperature overnight. Plates of phage serial dilutions exhibiting near complete lysis are then scraped with 5 – 10 mL of sterile 1× PBS (Invitrogen, Lithuania) and transferred to a 50 mL Falcon tube. The scrapings were vortexed vigorously followed by centrifugation at 5000 ×g for 10 minutes to pellet agar debris and the viral supernatant was sterilised with 0.45 µm syringe filters. The phage lysates were stored at 4°C for short-term purposes and at −80°C in 20% glycerol final concentration for long-term storage.

### Phage isolation from stream water

The water column in lotic systems functions as a dynamic conduit that integrates viral production from upstream biofilms through downstream dispersal, where waterborne phages represent a subset of mobile and infectious viruses dispersing across biofilm-associated hosts. Hence, concentrating viral particles from the water column enables scalable and reproducible recovery of viable phages. 120 L of fresh water was collected from a stream confluence located at Champéry, Switzerland that was fed by both groundwater-fed and glacier-fed streams (Fig.S1: 46.164 N, 6.8610 E; 1051 m above sea level). Water samples were immediately transported back to our facility and concentrated via tangential flow using a hollow fibre cartridge with 1.15 m^2^, 100 000 kDa membrane (GE Healthcare, USA) to a final volume 2 L which was filtered via 0.45 µm Sterivex (Merck, Germany) to exclude eukaryotic and most prokaryotic organisms but not viruses. The filtered water was further concentrated to 500 mL using a tangential flow cassette with 100 000 kDa (Sartorius, Germany). The concentrated water is then screened for phages against 40 previously isolated bacterial host strains via soft-agar overlay by mixing 500 µL of overnight host culture with 1 mL of concentrated water and 2 mL of 0.4% agar before casting on 1.5% agar media plates. Phage plaques identified were cored with sterile wide-bore pipette tips and resuspended vigorously in 100 µL of 1× PBS. Phages in PBS were then serially diluted and plated using soft agar overlay to perform a subsequent round of coring resulting in double-purified phage isolates. Purified phages were amplified in broth or soft-agar media to generate high viral titres (10^8^ – 10^10^ PFU/mL) for downstream analyses.

### Transmission electron microscopy and plaque morphology imaging

TEM was performed with negative stain. Each sample was adsorbed on a glow-discharged carbon-coated copper grid 400 mesh (EMS, Hatfield, PA, USA) washed with deionized water and stained with Uranyl Acetate 1% for 30 seconds. Observations was made using a Talos L120C electron microscope (Thermo Fisher, Hillsboro, USA) operated at 120 kV. Digital images were collected using a CMOS camera Ceta-S (Thermo Fisher, Hillsboro, USA) 4098 × 4098 pixels.), using a defocus range between −1.5 µm and −2.5 µm. Images of plaque morphologies were obtained on serially-diluted pure phage cultures on 0.4% soft-agar with a Scan 300 automatic colony counter (Interscience, France).

### Phage DNA extraction and long-read sequencing

1 mL of phage lysate at 10^8^ PFU/mL minimum, were initially treated with 10 U of DNase (Thermo Scientific, USA) with 1× DNase buffer (diluted from 10× stock comprising of 0.1 M Tris pH 7.5, 1 mM CaCl_2_ and 25 mM MgCl_2_) for 2 h at 37°C to eliminate bacterial DNA within the lysate. DNase was then inactivated by incubating at 75°C for 30 minutes. Removal of bacterial DNA was verified via 16S rRNA PCR (V3/V4 primers 341F: 5’-TCGTCGGCAGCGTCAGATGTGTATAAGAGACAGCCT ACGGGNGGCWGCAG-3’ and 785R: 5’- GTC TCGTGGGCTCGGAGATGTGTATAAGAGACA GGACTACHVGGGTATCTAATCC-3’) with the following cycling conditions: initial denaturation at 95°C for 3 minutes, 30 cycles at 95°C for 30 seconds, 55°C for 30 seconds, 72°C for 30 seconds, 5-minute final extension using 1 µL of template; and subsequently, gel electrophoresis with a ∼550 bp product if PCR-positive for 16S rRNA. PCR negative lysates were extracted for phage DNA using two methods. For most phage isolates, Norgen Biotek column-based phage extraction kit (Norgen Biotek, Canada) was used following manufacturer’s protocol including 20 µL Proteinase K (PanReac AppliChem, Switzerland) treatment and an extended incubation at 55°C for 1 h during viral lysis step. However, isolates that are recalcitrant to column-based extraction were extracted following a modified DNA isopropanol precipitation protocol. Here, 10% SDS (Fisher Scientific, UK) was added to the phage lysate at a final concentration of 0.5%, mixed well, and allowed to incubate at ambient room temperature for 30 minutes. Sodium acetate (Sigma-Aldrich, USA) solution was then added to the lysate to a final concentration of 0.3 M followed by 0.6 – 0.7 volume of ice-cold isopropanol (Merck, Germany). The mixture was mixed by inversion and immediately centrifuged at 15 000 ×g at 4°C for 30 minutes. The supernatant was discarded, and the DNA pellet was washed with 500 µL of freshly prepared 70% ethanol (Fisher Scientific, UK) and pelleted at 15 000 ×g at 4°C for 30 minutes, twice. Once residual ethanol from the final wash was dried, the pellets were resuspended in 50 µL TE buffer overnight at 4°C for complete DNA resuspension. Extracted DNA concentration, quality, and integrity was assessed using Qubit® Fluorometer with 1× dsDNA High Sensitivity kit (Invitrogen, USA), Nanodrop® (Scientific, USA), and Genomic TapeStation® (Agilent, USA) following manufacturers’ protocols, respectively. DNA library preparation and MinION long-read sequencing were performed using Native Barcoding Kit 24 V14 (Oxford Nanopore Technologies, UK) following manufacturer protocol, using short fragment buffer to maximise read preservation at final wash step prior to eluting the genomic library for sequencing. When long-read data were insufficient to assemble complete phage genomes, DNA sequencing was performed on the Illumina MiSeq i100 platform using the TruSeq™ DNA PCR-Free Library Prep Kit (Illumina, USA) with a random fragmentation (Covaris E220 focused ultrasonicator). We caution that certain phage isolates especially those infecting *P. cyclaminis*, remain notoriously recalcitrant to DNA extraction while the *Sphingomonas* phage isolate was limited to low amplification titre of <10^4^ PFU/mL despite our best efforts, which not only hampered DNA extraction but also transmission electron microscopy.

### Phage genome assembly, annotation, and bioinformatic analyses

Long-reads were base-called and demultiplexed using Dorado (v0.9.1) with high-accuracy model and remaining adapters and barcodes were removed using Porechop (v0.2.4). Any reads mapping to the bacterial host genome were also removed. Sequenced reads were assembled into contigs with Flye (v2.9.5) when using long-reads only and with Unicycler (v0.5.1) when performing hybrid assembly. Contigs were quality-checked to verify their viral origins using VIBRANT (v1.2.1) and geNomad (v1.9.0). Where possible, we also derived taxonomic information of our completed genomes with geNomad to assess the novelty of our viral collection against publicly available databases. We then complemented CheckV (v1.0.3) with an in-house read-mapping visualiser theBIGbam (v0.1; unpublished) to evaluate phage genome completeness based on nucleotide sequence and read mapping features, respectively. We dereplicated our collection of complete phage genomes via nucleotide identity using scripts provided by CheckV with 85% coverage and 95% nucleotide identity thresholds. Phage contigs assembled as head-to-tail concatemers were identified as overcomplete genomes, and genome lengths were automatically corrected by retaining a single-repeat unit corresponding to the expected complete phage genome. Genes were predicted using PHANOTATE (v1.5.0) and annotated with a combination of pharokka (v1.8.2) and phold (v1.1.0) using sequence and structural homology, respectively. DNA packaging mechanisms were also determined with theBIGbam using the logic from PhageTerm^33^. Nucleotide BLAST to NCBI database was performed with default settings using the dereplicated genome sequences as the query sequence. Meanwhile, queries to planetary-scale LOGAN contig database were initiated by randomly sampling 1 kb segments from the genomes of each dereplicated phage isolate. The 1 kb query sequence was then interrogated against all available contigs in LOGAN with the threshold setting set to the lowest value at 0.25 k-mer coverage (query date 01-Dec-2025). Cumulative k-mer coverage scores were calculated by summing the individual coverage values per source. All data was analysed and plotted with R (v.4.2.0).

### Crystal violet static biofilm assay

Bacterial overnight cultures were adjusted to OD_600_ ∼0.02 with fresh media and 190 µL was aliquoted into each well of a 96-well flat-bottom polystyrene plate (Greiner Bio-One, Switzerland). Biofilms were then grown statically in triplicate for each bacterial species for 24 h. Following static incubation, planktonic cells in the supernatant were discarded by carefully inverting the plate into a waste basin and the biofilms were wash thrice with MilliQ water (Merck Millipore, Germany) by submerging the plate in a separate filled basin and discarding the water in waste. The 96-well plate was then inverted on a paper towel to remove excess liquid and is allowed to air-dry. Crystal violet staining was performed by adding 200 µL of 0.01% Crystal violet solution (Sigma-Aldrich, USA) in each well and stained at ambient temperature with low-speed orbital shaking for 30 minutes. Excess Crystal violet was discarded, and the stained biofilms were washed thrice with MilliQ water and allowed to air-dry. 200 µL of 70% ethanol (Fisher Scientific, UK) was then added to each well and incubated at ambient room temperature for 30 minutes with low orbital shaking to solubilise the Crystal violet. Absorbance values at 590 nm were obtained (A_590_) using Biotek Synergy H1 plate reader (Agilent, USA) and the datapoints were blanked with A_590_ values from negative control wells containing sterile media.

### Microfluidic biofilm time-course phage infection

Polydimethylsiloxane (PDMS) microfluidic devices with channel dimension of 40 µm height, 250 µm width, and 1 mm length were purchased (Wunderlichips, Switzerland) and sterilised with 70% ethanol followed by UV radiation for 30 minutes. Microfluidic biofilms were established by first seeding the channels with 10 µL of overnight bacterial cultures adjusted to OD_600_ ∼0.1 in fresh media. Cells were allowed to attach under static conditions for 2 h at ambient room temperature. Devices were then connected a media-filled syringe via autoclave-sterile PTFE tubing (ID: 0.56 mm and OD: 1.07 mm, Fisher Scientific, Switzerland) with the outlet emptying into a waste Eppendorf tube. Media was infused continuously with a syringe pump (New Era Pump Systems, USA) at 0.1 µL/min flow rate and biofilms were grown for 2 h at ambient room temperature. Following overnight growth, sterile media syringes were swapped for 10^6^ PFU/mL of phages diluted in sterile media and phages were infused continuously at 0.1 µL/min flow rate for a further 96 h. Control biofilms not infected with phages were maintained with sterile media infusions. Biofilm development was then monitored over the 96-h duration under default brightfield settings via Zeiss Axio Zoom.V16 microscope with the Plan-NEOFLUAR Z 1.0×/0.25 FWD 56 mm objective and 112× zoom magnification (Carl Zeiss Microscopy, Germany). Images were acquired and processed with Zen (blue edition) software (Carl Zeiss Microscopy, Germany).

### Data, code, and phage culture availability

All code used for long-read preprocessing, assembly and genome annotation were integrated in a Snakemake (v7.32.4) pipeline available at: https://github.com/bhagavadgitadu22/PhageID. Sequencing reads and assembled genomes will be deposited in NCBI under BioProject: PRJNA1423127 following manuscript review. Purified phage cultures will also be submitted to CRBIP Biological Resource Center of Insitut Pasteur following manuscript acceptance. All plots were generated with R and raw data are available via supplementary data tables.

## Acknowledgements

We thank Martina Gonzalez and David Touchette for sharing their bacterial culture collection from streambed biofilms as bait for our phage isolation efforts. We also thank Florence Jagorel and Marc Monot from the group PhagoMics, Phages.fr, for their advice and support, and the Biomics Platform, C2RT, Institut Pasteur, Paris, France, supported by France Génomique (ANR-10-INBS-09) and IBISA. We finally thank Tom Battin for his guidance and provision of laboratory facility in support of this work. This work was supported by the Swiss National Science Foundation under grant no.212726 awarded to Hannes Peter.

## Author contributions

W.H.C and H.P. conceived the study, and W.H.C designed experiments and interpreted data. W.H.C, M.B., and A.H. carried out field expeditions, sample processing, and phage isolations. W.H.C and A.H. purified phages and maintained the phage collection throughout the study. W.H.C, A.H., and F.B. performed phage genome extractions and sequencing. M.B. established bioinformatic pipelines and analysed genomes. W.H.C performed experiments, viral database screens, and data analyses. D.D. performed electron microscopy imaging. W.H.C prepared figures. W.H.C. and H.P. wrote the manuscript. H.P. supervised the study. All authors reviewed, edited, and commented on the manuscript.

## Notes

https://www.ncbi.nlm.nih.gov/search/all/?term=PRJNA1423127

### Competing Interest Statement

The authors have declared no competing interest.

## References

1. Flemming, H. C. & Wuertz, S. Bacteria and archaea on Earth and their abundance in biofilms. Nature Reviews Microbiology 2019 17:4 17, 247–260 (2019).

2. Flemming, H. C. et al. Biofilms: an emergent form of bacterial life. Nature Reviews Microbiology 2016 14:9 14, 563–575 (2016).

3. Battin, T. J., Besemer, K., Bengtsson, M. M., Romani, A. M. & Packmann, A. I. The ecology and biogeochemistry of stream biofilms. Nature Reviews Microbiology 2016 14:4 14, 251–263 (2016).

4. Thorpe, A. C. et al. National-scale biogeography and function of river and stream bacterial biofilm communities. Nature Communications 2025 16:1 16, 10571- (2025).

5. Michoud, G. et al. Unexpected functional diversity of stream biofilms within and across proglacial floodplains despite close spatial proximity. Limnol. Oceanogr. 68, 2183–2194 (2023).

6. Bourquin, M. et al. Predicting climate-change impacts on the global glacier-fed stream microbiome. Nature Communications 2025 16:1 16, 1264- (2025).

7. Hall-Stoodley, L., Costerton, J. W. & Stoodley, P. Bacterial biofilms: from the Natural environment to infectious diseases. Nature Reviews Microbiology 2004 2:2 2, 95–108 (2004).

8. Chevallereau, A., Pons, B. J., van Houte, S. & Westra, E. R. Interactions between bacterial and phage communities in natural environments. Nature Reviews Microbiology 2021 20:1 20, 49–62 (2021).

9. Hu, C. et al. To kill or to piggyback: Switching of viral lysis-lysogeny strategies depending on host dynamics. Science of The Total Environment 959, 178233 (2025).

10. Piel, D. et al. Phage–host coevolution in natural populations. Nature Microbiology 2022 7:7 7, 1075–1086 (2022).

11. Borin, J. M. et al. Rapid bacteria-phage coevolution drives the emergence of multiscale networks. Science (1979). 382, 674–678 (2023).

12. Shiah, F. K. et al. Viral shunt in tropical oligotrophic ocean. Sci. Adv. 8, 2829 (2022).

13. Gilbert, N. E. et al. Seasonal enhancement of the viral shunt catalyzes a subsurface oxygen maximum in the Sargasso Sea. Nature Communications 2025 17:1 17, 352-(2025).

14. Rosenwasser, S., Ziv, C., Creveld, S. G. van & Vardi, A. Virocell Metabolism: Metabolic Innovations During Host–Virus Interactions in the Ocean. Trends Microbiol. 24, 821–832 (2016).

15. Howard-Varona, C. et al. Phage-specific metabolic reprogramming of virocells. The ISME Journal 2020 14:4 14, 881–895 (2020).

16. Pires, D. P., Melo, L. D. R. & Azeredo, J. Understanding the Complex Phage-Host Interactions in Biofilm Communities. Annu. Rev. Virol. 8, 73–94 (2021).

17. Vidakovic, L., Singh, P. K., Hartmann, R., Nadell, C. D. & Drescher, K. Dynamic biofilm architecture confers individual and collective mechanisms of viral protection. Nature Microbiology 2017 3:1 3, 26–31 (2017).

18. Winans, J. B., Zeng, L. & Nadell, C. D. Spatial propagation of temperate phages within and among biofilms. Proc. Natl. Acad. Sci. U. S. A. 122, e2417058122 (2025).

19. Prentice, J. A., van de Weerd, R. & Bridges, A. A. Cell-lysis sensing drives biofilm formation in Vibrio cholerae. Nature Communications 2024 15:1 15, 2018- (2024).

20. Vaidya, S. et al. Bacteria use exogenous peptidoglycan as a danger signal to trigger biofilm formation. Nature Microbiology 2025 10:1 10, 144–157 (2025).

21. Winans, J. B., Wucher, B. R. & Nadell, C. D. Multispecies biofilm architecture determines bacterial exposure to phages. PLoS Biol. 20, e3001913 (2022).

22. Gliźniewicz, M. et al. Advances in bacteriophage-mediated strategies for combating polymicrobial biofilms. Front. Microbiol. 14, 1320345 (2023).

23. Bekliz, M., Pramateftaki, P., Battin, T. J. & Peter, H. Viral diversity is linked to bacterial community composition in alpine stream biofilms. ISME Communications 2, (2022).

24. Michoud, G. et al. Mapping the metagenomic diversity of the multi-kingdom glacier-fed stream microbiome. Nature Microbiology 2024 10:1 10, 217–230 (2025).

25. Resch, G. et al. Between Centralization and Fragmentation: The Past, Present, and Future of Phage Collections. PHAGE: Therapy, Applications, and Research 5, 22 (2024).

26. Russell, D. A. & Hatfull, G. F. PhagesDB: the actinobacteriophage database. Bioinformatics 33, 784–786 (2017).

27. Maffei, E. et al. Systematic exploration of Escherichia coli phage–host interactions with the BASEL phage collection. PLoS Biol. 19, e3001424 (2021).

28. Fletcher, J. et al. The Citizen Phage Library: Rapid Isolation of Phages for the Treatment of Antibiotic Resistant Infections in the UK. Microorganisms 12, 253 (2024).

29. Touchette, D. et al. Experimental evidence on the impact of climate-induced hydrological and thermal variations on glacier-fed stream biofilms. FEMS Microbiol. Ecol. 101, 163 (2025).

30. Ezzat, L. et al. Diversity and biogeography of the bacterial microbiome in glacier-fed streams. Nature 2025 637:8046 637, 622–630 (2025).

31. Kohler, T. J. et al. Global emergent responses of stream microbial metabolism to glacier shrinkage. Nature Geoscience 2024 17:4 17, 309–315 (2024).

32. Harding, K. R., Kyte, N. & Fineran, P. C. Jumbo phages. Current Biology 33, R750–R751 (2023).

33. Valdez, A., Sun, H., Weiss, H. H. & Aranson, I. Biomechanical modeling of spatiotemporal bacteria-phage competition. Communications Physics 2025 8:1 8, 139- (2025).

34. Knecht, L. E., Veljkovic, M. & Fieseler, L. Diversity and Function of Phage Encoded Depolymerases. Front. Microbiol. 10, 502215 (2020).

35. Oliveira, H. et al. Ability of phages to infect Acinetobacter calcoaceticus-Acinetobacter baumannii complex species through acquisition of different pectate lyase depolymerase domains. Environ. Microbiol. 19, 5060–5077 (2017).

36. Qin, K. et al. Isolation and characterization of wetland VSW-3, a novel lytic cold-active bacteriophage of Pseudomonas fluorescens. Can. J. Microbiol. 63, 110–118 (2017).

37. Gittrich, M. R. et al. Isolation and characterization of 24 phages infecting the plant growth-promoting rhizobacterium Klebsiella sp. M5al. PLoS One 20, e0313947 (2025).

38. Qin, X. et al. Psychrophilic phage phiGM22-3 efficiently controls Pseudomonas fluorescens contamination in cold-stored milk. Int. J. Food Microbiol. 411, 110525 (2024).

39. Chikhi, R. et al. Logan: Planetary-Scale Genome Assembly Surveys Life’s Diversity. bioRxiv 2024.07.30.605881 (2025) doi:10.1101/2024.07.30.605881.

40. Chaudhari, H. V., Inamdar, M. M. & Kondabagil, K. Scaling relation between genome length and particle size of viruses provides insights into viral life history. iScience 24, 102452 (2021).

41. Garneau, J. R., Depardieu, F., Fortier, L. C., Bikard, D. & Monot, M. PhageTerm: a tool for fast and accurate determination of phage termini and packaging mechanism using next-generation sequencing data. Scientific Reports 2017 7:1 7, 8292- (2017).

42. Yang, J. Y. et al. Degradation of host translational machinery drives tRNA acquisition in viruses. Cell Syst. 12, 771–779.e5 (2021).

43. Encina-Robles, J., Pérez-Villalobos, V. & Bustamante, P. The HicAB System: Characteristics and Biological Roles of an Underappreciated Toxin-Antitoxin System. International Journal of Molecular Sciences 2024, Vol. 25, Page 12165 25, 12165 (2024).

44. Martinez, S. & Hausinger, R. P. Catalytic Mechanisms of Fe(II)- and 2-Oxoglutarate-dependent Oxygenases. Journal of Biological Chemistry 290, 20702–20711 (2015).

45. Mahler, M. et al. Phage arabinosyl-hydroxy-cytosine DNA modifications result in distinct evasion and sensitivity responses to phage defense systems. Cell Host Microbe 33, 1173–1190.e9 (2025).

46. Richard, H., Carpenter, E. J., Komada, T., Palmer, P. T. & Rochman, C. M. Biofilm facilitates metal accumulation onto microplastics in estuarine waters. Science of The Total Environment 683, 600–608 (2019).

47. Goff, J. L. et al. Tellurite Adsorption onto Bacterial Surfaces. Environ. Sci. Technol. 55, 10378–10386 (2021).

48. Boles, B. R. & Singh, P. K. Endogenous oxidative stress produces diversity and adaptability in biofilm communities. Proc. Natl. Acad. Sci. U. S. A. 105, 12503–12508 (2008).

49. Chiang, B. Y. et al. Protein S-thiolation by glutathionylspermidine (Gsp): The role of Escherichia coli Gsp synthetase/amidase in redox regulation. Journal of Biological Chemistry 285, 25345–25353 (2010).

50. Hutinet, G. et al. 7-Deazaguanine modifications protect phage DNA from host restriction systems. Nature Communications 2019 10:1 10, 5442- (2019).

51. Díaz-Rullo, J. & González-Pastor, J. E. tRNA queuosine modification is involved in biofilm formation and virulence in bacteria. Nucleic Acids Res. 51, 9821–9837 (2023).

52. Bailey, Z. M., Igler, C. & Wendling, C. C. Prophage maintenance is determined by environment-dependent selective sweeps rather than mutational availability. Current Biology 34, 1739–1749.e7 (2024).

53. Simmons, E. L. et al. Biofilm Structure Promotes Coexistence of Phage-Resistant and Phage-Susceptible Bacteria. mSystems 5, (2020).

54. Bond, M. C., Vidakovic, L., Singh, P. K., Drescher, K. & Nadell, C. D. Matrix-trapped viruses can prevent invasion of bacterial biofilms by colonizing cells. Elife 10, e65355 (2021).

55. Magar, S. et al. Characterization and genomic analysis of Sharanji: a jumbo bacteriophage of Escherichia coli. Virol. J. 22, 67 (2025).

56. Peter, H., Michoud, G., Busi, S. B. & Battin, T. J. The role of phages for microdiverse bacterial communities in proglacial stream biofilms. Frontiers in Microbiomes 2, 1279550 (2023).

57. Breitbart, M., Bonnain, C., Malki, K. & Sawaya, N. A. Phage puppet masters of the marine microbial realm. Nature Microbiology 2018 3:7 3, 754–766 (2018).

58. Immerzeel, W. W. et al. Importance and vulnerability of the world’s water towers. Nature 2019 577:7790 577, 364–369 (2019).

59. Rose, R., Constantinides, B., Tapinos, A., Robertson, D. L. & Prosperi, M. Challenges in the analysis of viral metagenomes. Virus Evol. 2, (2016).

60. Bekliz, M., Brandani, J., Bourquin, M., Battin, T. J. & Peter, H. Benchmarking protocols for the metagenomic analysis of stream biofilm viromes. PeerJ 2019, e8187 (2019).

61. Subedi, D. & Barr, J. J. Temporal Stability and Genetic Diversity of 48-Year-Old T-Series Phages. mSystems 6, (2021).

62. Zhong, Z. P. et al. Glacier-preserved Tibetan Plateau viral community probably linked to warm–cold climate variations. Nature Geoscience 2024 17:9 17, 912–919 (2024).

63. Luque, A., Benler, S., Lee, D. Y., Brown, C. & White, S. The Missing Tailed Phages: Prediction of Small Capsid Candidates. Microorganisms 2020, Vol. 8, Page 1944 8, 1944 (2020).

64. Dunsing, V., Irmscher, T., Barbirz, S. & Chiantia, S. Purely Polysaccharide-Based Biofilm Matrix Provides Size-Selective Diffusion Barriers for Nanoparticles and Bacteriophages. Biomacromolecules 20, 3842–3854 (2019).

65. Comeau, A. M., Bertrand, C., Letarov, A., Tétart, F. & Krisch, H. M. Modular architecture of the T4 phage superfamily: A conserved core genome and a plastic periphery. Virology 362, 384–396 (2007).

66. Bouras, G. et al. Protein structure-informed bacteriophage genome annotation with Phold. Nucleic Acids Res. 54, (2026).

67. Jo, J., Price-Whelan, A. & Dietrich, L. E. P. Gradients and consequences of heterogeneity in biofilms. Nature Reviews Microbiology 2022 20:10 20, 593–607 (2022).

68. Wilde, J. et al. Assessing phage-host population dynamics by reintroducing virulent viruses to synthetic microbiomes. Cell Host Microbe 32, 768–778.e9 (2024).

69. Castledine, M., Padfield, D., Schoeman, M., Berry, A. & Buckling, A. Bacteria–phage (co)evolution is constrained in a synthetic community across multiple bacteria–phage pairs. Microbiology (United Kingdom) 171, 001577 (2025).

70. Depetris, A. et al. Biophysical properties at patch scale shape the metabolism of biofilm landscapes. npj Biofilms and Microbiomes 2022 8:1 8, 5- (2022).

71. Carrara, F., Altermatt, F., Rodriguez-Iturbe, I. & Rinaldo, A. Dendritic connectivity controls biodiversity patterns in experimental metacommunities. Proc. Natl. Acad. Sci. U. S. A. 109, 5761–5766 (2012).

