## Supplementary Figures S1 - S2 & Tables S1 - S2 for "Viral isolation reveals novel and diverse phages infecting natural stream biofilms"

### Supplementary information (SI)

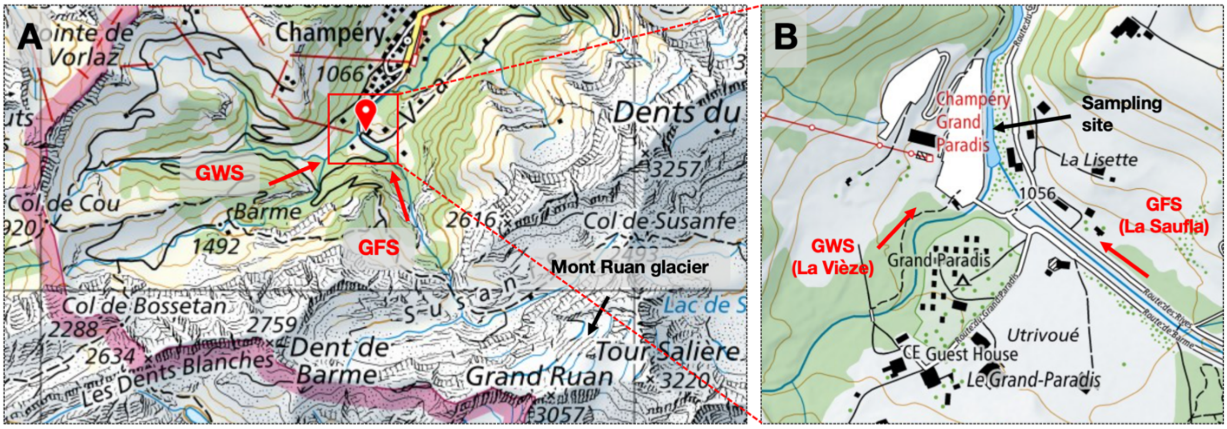


**Fig.S1 Location of stream water sampling site.** A) Elevated map view of sampling site indicated by red positional indicator which sits at a confluence of two alpine streams: a groundwater-fed stream (GWS) from the direction of Col de Cou high mountain pass and a glacier-fed stream (GFS) originating from the Mont Ruan glacier. B) View of the map inset demonstrating the exact sampling site at the stream confluence from the La Vièze GWS and La Saufla GFS. Exact coordinates and elevation from sea level of the sampling site are as follows: 46.164 N, 6.8610 E; 1051 m. Map layout acquired via the Swiss Federal Office of Topography (Swisstopo).

**Tail fibre amino acid sequence for phage isolates B503P1 and B503P2**

```
                     > Receptor binding domain (homology with T7 phage tail fibre)
B503P1   1   MAINTIKLYELNGVVTRFPVTFEYLARKFVKVTLLGDERTELTLLTDFVFEGSAGIRLLG   60
             ||||||||||||||||||||||||||||||||||||||||||||||||||||||||||||
B503P2   1   MAINTIKLYELNGVVTRFPVTFEYLARKFVKVTLLGDERTELTLLTDFVFEGSAGIRLLG   60

B503P1  61   TWGGGDGYEQVEIRRETSASERIVSFSDATILRAGDLNTAELQSLHIAEEARDQTSQALS   120
             ||||||||||||||||||||||||||||||||||||||||||||||||||||||||||||
B503P2  61   TWGGGDGYEQVEIRRETSASERIVSFSDATILRAGDLNTAELQSLHIAEEARDQTSQALS   120

B503P1 121   RDEDGNIDAQIHRVVNMEDGINPQDGVTLHQLQEATSENATTLRNDLIGPDGGKMVSIRP   180
             ||||||||||||||||||||||||||||||||||||||||||||||||||||||||||||
B503P2 121   RDEDGNIDAQIHRVVNMEDGINPQDGVTLHQLQEATSENATTLRNDLIGPDGGKMVSIRP   180

B503P1 181   AYSGAIAQSLESILGQTLNVTNFGVHPSNSAAANTVGMNKANLEAVRLGLPLFFPPIGVY   240
             ||||||||||||||||||||||||||||||||||||||||||||||||||||||||||||
B503P2 181   AYSGAIAQSLESILGQTLNVTNFGVHPSNSAAANTVGMNKANLEAVRLGLPLFFPPIGVY   240

B503P1 241   MMGRLADVTGKLTWYSHSTHAVIQVGGDVWPYDVSPAPFLTVYGSWAAHSLGLDQNWVTP   300
             ||||||||||||||||||||||||||||||||||||||||||||||||||||||||||||
B503P2 241   MMGRLADVTGKLTWYSHSTHAVIQVGGDVWPYDVSPAPFLTVYGSWAAHSLGLDQNWVTP   300

B503P1 301   LHGNVSGPYRDDMNPRTWGGNWMVKVVATTKDSLLVTDCALKTVARGFYAEALDPALGGA   360
             ||||||||||||||||||||||||||||||||||||||||||||||||||||||||||||
B503P2 301   LHGNVSGPYRDDMNPRTWGGNWMVKVVATTKDSLLVTDCALKTVARGFYAEALDPALGGA   360

B503P1 361   DSVVMNGCTGDNFSTASQCMIAADSPGSFQAIHNRIATTKWTDGETYMGNGTSVVFPFGG   420
             ||||||||||||||||||||||||||||||||||||||||||||||||||||||||||||
B503P2 361   DSVVMNGCTGDNFSTASQCMIAADSPGSFQAIHNRIATTKWTDGETYMGNGTSVVFPFGG   420

B503P1 421   TNIQILDNHFSGYQLVARGPSSNPWVPNTYFGPDMEFRTPSGEARRVTTAYTTGPVYGLL   480
             ||||||||||||||||||||||||||||||||||||||||||||||||||||||||||||
B503P2 421   TNIQILDNHFSGYQLVARGPSSNPWVPNTYFGPDMEFRTPSGEARRVTTAYTTGPVYGLL   480
                                                > Pectin lyase β-helix (partial homology)
B503P1 481   DVANTLLIYNPQQKLHICRNTILAPIADTAAYGWDECIINENFIEDSGDMGIATSGSGFL   540
             ||||||||||||||||||||||||||||||||||||||||||||||||||||||||||||
B503P2 481   DVANTLLIYNPQQKLHICRNTILAPIADTAAYGWDECIINENFIEDSGDMGIATSGSGFL   540
                                                S590N substitution: v
B503P1 541   CVCDNIINRTRNGGIDVTYGGIINVAGNIVKDYARASGGVYKRIDTINPSVWASNGGFNL   600
             ||||||||||||||||||||||||||||||||||||||||||||||||| ||||||||||
B503P2 541   CVCDNIINRTRNGGIDVTYGGIINVAGNIVKDYARASGGVYKRIDTINPNVWASNGGFNL   600

B503P1 601   AGITVGLTQEPGTKQVNITGNNLYFTSLPPISDDGPNNGGIVRRAVHGIYSQVSSNVNAQ   660
             ||||||||||||||||||||||||||||||||||||||||||||||||||||||||||||
B503P2 601   AGITVGLTQEPGTKQVNITGNNLYFTSLPPISDDGPNNGGIVRRAVHGIYSQVSSNVNAQ   660

B503P1 661   NTINATGNMIQDNQADMPNFYLAVATLRFYNKAYTGEIVPGGLYKDGATQFVLVGFLGEG   720
             ||||||||||||||||||||||||||||||||||||||||||||||||||||||||||||
B503P2 661   NTINATGNMIQDNQADMPNFYLAVATLRFYNKAYTGEIVPGGLYKDGATQFVLVGFLGEG   720

B503P1 721   GLVFVKQFRGAASPFPGQVFVGPNGSTITTADNPQLGFIGSNESGNVDFTTKYSPDNYEL   780
             ||||||||||||||||||||||||||||||||||||||||||||||||||||||||||||
B503P2 721   GLVFVKQFRGAASPFPGQVFVGPNGSTITTADNPQLGFIGSNESGNVDFTTKYSPDNYEL   780

B503P1 781   VYRGAEMVTLAPIPTGTFKSFEATVAGAGFYDFCQATAPYQLQGLEVTSYVSGANKVTTI   840
             ||||||||||||||||||||||||||||||||||||||||||||||||||||||||||||
B503P2 781   VYRGAEMVTLAPIPTGTFKSFEATVAGAGFYDFCQATAPYQLQGLEVTSYVSGANKVTTI   840

B503P1 841   VKNSTGSTVTLEPAVWRLKVERF   863
             |||||||||||||||||||||||
B503P2 841   VKNSTGSTVTLEPAVWRLKVERF   863
```

**Fig.S2 Non-synonymous mutation of tail fiber protein of Janthinobacterium phage B503P2.** Bolded text indicates the polar amino acid substitution from serine to asparagine at position 590 (S590N). Highlights indicate regions demonstrating amino acid sequence homology to known protein motifs via MOTIF search (https://www.genome.jp/tools/motif/) against Pfam, NCBI-Conserved Domain Database (CDD), and Expasy PROSITE databases using default settings. Yellow region corresponds to homology with the receptor binding domains of T7 phage tail fibre[1] [NCBI-CDD ID: 222790, Score: 172, E-value: 2e-45, Date: 11 Feb 2026] while blue region exhibited partial homology with a right-handed β-helix associated with pectate lyase [NCBI-CDD ID: 463811, Score: 32.4, E-value: 0.56, Date: 11 Feb 2026], where the latter was also reported in phages as depolymerases to degrade bacterial capsule[2].

**Table S1: List of 37 alpine bacterial isolates and total number of phage isolated**

| Isolate # | Taxonomic classification | Total # of phages isolated (non-dereplicated) | Total successful sequenced |
|---|---|---|---|
| 1 | *Iodobacter* sp. | 0 | n/a |
| 2 | *Chryseobacterium aquaticum* | 0 | n/a |
| 3 | *Arthrobacter sunyaminii* | 0 | n/a |
| 4 | *Exiguobacterium antarticum* | 0 | n/a |
| 5 | *Aeromonas salmonicida* | 0 | n/a |
| 6 | *Microbacterium* sp. | 0 | n/a |
| 7 | *Comamonas jiangduensis* | 0 | n/a |
| 8 | *Duganella* sp. | **1** | **1** |
| 9 | *Pseudarthrobacter sulfonivorans* | 0 | n/a |
| 10 | *Janthinobacterium* sp.002735705 | **1** | **1** |
| 11 | *Comamonas acidovorans* | **1** | **1** |
| 12 | *Acinetobacter guillouiae* | 0 | n/a |
| 13 | *Yersinia kristensenii* | 0 | n/a |
| 14 | *Arthrobacter* sp. | 0 | n/a |
| 15 | *Chryseobacterium* sp. | 0 | n/a |
| 16 | *Deinococcus* sp. | 0 | n/a |
| 17 | *Rahnella inusitata* | **22** | **18** |
| 18 | *Massilia* sp. | **5** | **3** |
| 19 | *Acinetobacter* sp. | 0 | n/a |
| 20 | *Klebsiella terrigena* | 0 | n/a |
| 21 | *Pedobacter* sp. | 0 | n/a |
| 22 | *Pseudomonas* sp. | **2** | **2** |
| 23 | *Zooglea ramigera* | 0 | n/a |
| 24 | *Pseudomonas germanica* | 0 | n/a |
| 25 | *Acinetobacter bohemicus* | 0 | n/a |
| 26 | *Flavobacterium circumlabens* | 0 | n/a |
| 27 | *Pseudomonas cyclaminis* | **7** | **1** |
| 28 | *Pseudomonas fragi* | 0 | n/a |
| 29 | *Pseudomonas haemolytica* | **6** | **5** |
| 30 | *Flavobacterium* sp. | **3** | **3** |
| 31 | *Brevundimonas vesicularis* | **2** | **2** |
| 32 | *Flavobacterium brachiicola* | 0 | n/a |
| 33 | *Janthinobacterium tructae* | **2** | **2** |
| 34 | *Pseudomonas fluorescens* | **1** | **1** |
| 35 | *Rhodoferax* sp. 002943465 | **3** | **3** |
| 36 | *Sphingomonas* sp. | **1** | **0** |
| 37 | *Exiguobacterium undae* | 0 | n/a |
| | **Total:** | 57 | 43 |

NB Cells in red indicate cases where some phages were recalcitrant to DNA extractions

### Table S2: List of 30 unique phage isolates

| | Phage (φ) isolate | Genome extracted (Y/N?) | TEM image (Y/N?) | Number of re-isolations | Notes |
|---|---|---|---|---|---|
| 1 | Brevundimonas φ B307P1 | Y | Y | 0 | |
| 2 | Brevundimonas φ B307P2 | Y | N | 0 | |
| 3 | Comamonas jumboφ B146P1 | Y | Y | 0 | |
| 4 | Duganella φ B044P1 | Y | Y | 0 | |
| 5 | Flavobacterium φ B147P1 | Y | Y | 0 | |
| 6 | Flavobacterium φ B147P2 | Y | Y | 0 | |
| 7 | Janthinobacterium φ B057P1 | Y | Y | 0 | |
| 8 | Janthinobacterium φ B503P1 | Y | Y | 1 | • TEM including genome identical B503P2 isolate<br>• “Reisolation” includes the B503P2 isolate. |
| 9 | Massilia φ B343P1 | Y | Y | 0 | |
| 10 | Massilia φ B343P2 | Y | Y | 0 | |
| 11 | Massilia φ B343P3 | Y | N | 0 | |
| 12 | Pseudomonas φ B072P1 | Y | Y | 0 | |
| 13 | Pseudomonas φ B072P2 | Y | N | 3 | |
| 14 | Pseudomonas φ B427P1 | Y | Y | 1 | |
| 15 | Pseudomonas φ B508P1 | Y | N | 0 | |
| 16 | Pseudomonas φ B529P1 | Y | N | 0 | |
| 17 | Rahnella jumboφ B311P2 | Y | Y | 0 | |
| 18 | Rahnella proφ B311P3 | Y | N | 2 | |
| 19 | Rahnella φ B311P1 | Y | Y | 8 | |
| 20 | Rahnella φ B311P4 | Y | N | 0 | |
| 21 | Rahnella φ B311P5 | Y | N | 1 | |
| 22 | Rahnella φ B311P6 | Y | N | 0 | |
| 23 | Rahnella φ B311P7 | Y | N | 1 | |
| 24 | Rahnella φ B311P8 | Y | N | 1 | |
| 25 | Rahnella φ B311P9 | Y | Y | 0 | |
| 26 | Rhodoferax φ B534P1 | Y | Y | 0 | |
| 27 | Rhodoferax φ B534P2 | Y | Y | 0 | |
| 28 | Rhodoferax φ B534P3 | Y | Y | 0 | |

### SI references

1. Garcia-Doval, C. & Van Raaij, M. J. Structure of the receptor-binding carboxy-terminal domain of bacteriophage T7 tail fibers. *Proc. Natl. Acad. Sci. U. S. A.* 109, 9390–9395 (2012).
2. Oliveira, H. *et al.* Ability of phages to infect Acinetobacter calcoaceticus-Acinetobacter baumannii complex species through acquisition of different pectate lyase depolymerase domains. *Environ. Microbiol.* 19, 5060–5077 (2017).
